# Nol12 is a multifunctional endonuclease at the nexus of RNA and DNA metabolism

**DOI:** 10.1101/043935

**Authors:** D. D. Scott, C. Trahan, P.J. Zindy, L.C. Aguilar, M.Y. Delubac, K. E. Wei, M. Oeffinger

**Author notes:** The authors wish it to be known that, in their opinion, the first three authors should be regarded as joint First Authors. Current Address: Human Oncology and Pathogenesis Program, Memorial Sloan Kettering Cancer Center, New York, New York (K. E. Wei).

## Abstract

Endo‐ and exonucleases are major contributors to RNA metabolism through their diverse roles in maturation and turnover of different species of RNA as well as transcription. Recent data suggests RNA nucleases also affect genome stability programs and act along DNA repair pathways. Here, we describe Nol12 as a multifunctional RNA/DNA endonuclease found in different subcellular compartments - the nucleoplasm, where it co-localizes with the RNA/DNA helicase Dhx9 and paraspeckles, nucleoli as well as GW/P-bodies. We show that Nol12 is required for a key step in ribosomal RNA processing, separating large and small subunit precursors at site 2, rerouting ribosome biogenesis via an alternative pathway in its absence to ensure ribosome production. Furthermore, loss of Nol12 results in increased oxidized DNA levels followed by a rapid p53-independent ATR-Chk1-mediated apoptotic response, suggesting a role for Nol12 in the prevention or resolution of oxidative DNA damage. Identification of a complex Nol12 interactome, which includes NONO, Dhx9 and DNA-PK, further supports its diverse functions in RNA metabolism and DNA maintenance, establishing Nol12 as a multifunctional endonuclease.

## INTRODUCTION

Cells contain many different species of RNA, all of which undergo extensive processing to either achieve their mature functional form, or to be degraded as part of quality control or turnover mechanisms (1, 2). Endo‐ and exonucleases are the essential players along all of these pathways, and, in most cases, they act in concert on specific RNA substrates. However, many nucleases have multiple functions and are not just acting on one single pathway or species of RNA, but are rather involved in the processing and/or turnover of different types of RNAs along different pathways. Such multifunctionality has been demonstrated for several endo and exonucleases in higher eukaryotes. For example, the nuclear 5’-3’ exonuclease Xrn2 has a wide range of conserved cellular functions on a variety of RNA species including processing of pre-rRNA, nuclear RNA quality control, transcription termination, snoRNA maturation and miRNA turnover (3). A number of components of the multi-nuclease exosome complex were also shown to function on diverse RNA substrates within different cellular subcompartments, with hRrp6/PM/Scl-100 fulfilling several roles in nuclear RNA metabolism, including nucleolar pre-rRNA processing and nuclear mRNA surveillance (4, 5), while hDis3 was found to act in surveillance as well as degradation of aberrant pre-mRNAs and pre-snoRNAs (6, 7) and has also been implicated in paraspeckles formation (6). Besides exonucleases, recent studies have also highlighted the roles of endonucleases in RNA metabolism in higher eukaryotes, and several were shown to contribute to RNA turnover in various subcellular compartments (8). Amongst them hDis3, which contains dual exo-endonuclease activity, degrading both aberrant rRNA processing precursor and cryptic unstable transcripts (CUTs) via its endonuclease activity in the nucleus, while its variant, hDis3L, functions exonucleolytically in the cytoplasm (9-11). Another endonuclease, Rnt1/RNaseIII, was reported to process both pre-rRNAs and snoRNAs, whereas multiple endonucleases, Dicer, Ago2 and C3PO, are involved in the maturation of small interfering RNAs; in addition, Dicer also processes pre-micro RNAs into their mature forms (8).

Besides their numerous and often diverse roles in RNA metabolism, mounting evidence over the last few years supports the idea that RNA-binding and processing proteins are not only involved in different steps of RNA life but can also affect genome stability programs and even function directly in DNA repair. A number of large-scale genetic and proteomic studies of proteins involved in DNA DDR showed enrichment for RNA processing proteins, suggesting that RNA metabolism and DNA repair pathways intersect functionally (12). Along such lines, requirement for Rrp6 in the repair of DNA doublestrand breaks (DSBs) has recently been demonstrated in HeLa cells (13), and Xrn2 was show to cooperate with the helicase senataxin to resolve RNA:DNA hybrids (14). A function for the endonuclease Dicer in DNA damage and repair has also been shown (15, 16). Moreover, known RNA-protein structures within the nucleus, such as paraspeckles and the nucleolus, and some of their components have been implicated in DNA damage response (DDR) (17, 18). While the exact roles played by these nucleases in the cell’s response to different sources of DNA damage is still mostly unexplored, different mechanisms for such cross-functionality have been proposed (19). Involvement on the level of mRNA regulation, direct participation in the DDR, or prevention of DNA damage are all suggested modes of function of how RNA processing proteins could impact the maintenance of genome integrity.

In this manuscript we describe the role of the human protein Nol12 in both RNA and DNA metabolism. Previously, the *Drosophila* Nol12-homologue *Viriato* was shown to modulate signaling during eye development, and its interactor genes were overwhelmingly involved in development of the nervous system (20). Loss of *Viriato* resulted in cell proliferation, developmental delay and apoptosis. Moreover, *Viriato* as well as the mouse Nol12-homologue Nop25 were shown to affect nucleolar integrity (20, 21), while its yeast homologue, Rrp17, was identified as a putative 5’-3’ exonuclease with function in ribosome biogenesis (22); a potential role for Nol12 in ribosome maturation has also been suggested (4). However, public human proteome data sets analyzing protein profiles showed localization of human Nol12 not only within the nucleolus but also the nucleo‐ and cytoplasm (23), suggesting functions for Nol12 outside of the nucleolus and ribosome maturation. In line with this, Nol12 was previously identified to interact with p33Monox, a human neuron-expressed cytosolic protein and inhibitor of amyloid beta precursors protein (APP) and Bcl2 phosphorylation from a human brain cDNA library (24).

Here, we show Nol12 to be a multifunctional nuclease exhibiting endonucleolytic activity *in vitro* on RNA and DNA substrates. The identification of a diverse Nol12 interactome, supported by immunofluorescence microscopy, suggests a role for the nuclease in RNA metabolism in different subcellular compartments such as nucleoli, the paraspeckles, and P-bodies. Nol12 is required for a key-processing step in ribosomal RNA processing, separating small and large subunit rRNAs. In addition, we show that Nol12 has a role in management of oxidized DNA that is linked to a rapid pro-apoptotic ATR-mediated DNA damage response in its absence. Moreover, Nol12 associates and co-localizes with the DNA/RNA DEAD-box helicase Dh×9 as well as paraspeckle components NONO and SfpQ, further indicating a role for Nol12 in genome instability prevention. Together with our identification of a complex Nol12-interactome, these data suggest a function for the protein along various RNA metabolism pathways and DNA maintenance processes, providing insights into the diverse roles of an endonuclease at the intersection between RNA - DNA life and surveillance.

## MATERIALS AND METHODS

### DNA constructs

For recombinant Nol12 (rNol12) expression, the cDNA of NOL12 was amplified by PCR from pRS414-3×HA-NOL12 (22) with oligonucleotides NdeI-NOL12-F (5’-ccc<u>catatgg</u>gccgcaacaagaagaagaa-3’) and XhoI-NOL12-R (5’-ggg<u>ctcgag</u>cgggccctgaaacagcacttccaggccgctgctctccccgctgtgccgtgc-3’). The PCR product was cloned into NdeI-XhoI sites of pET21a to generate plasmid pET21a-NOL12-6xHis. In order to construct a PrA-NOL12 encoding plasmid for Flip-in recombination Flp-In T-REx system (see cell culture section), the ORF of NOL12 was amplified by PCR from the same template as above with oligonucleotides NcoI-NOL12-F (5’-ggaatcaggaccatgggaaccggccgcaacaagaagaagaagcgagatggtga-3’) and NotI-NOL12-R (5’-ggaatcaggagcggccgcctccccgctgtgccgtgctttgcctgtgagacggcggcgct-3’). The amplicon was cloned into NcoI-NotI sites of pENTR4 to generate pENTR4-NOL12. Protein A was amplified by PCR from pBXA (25) with oligonucleotides HindIII-PrA-F (5’-gctagc<u>aagcttg</u>aattcatggttggtactttctatcg-3’) and HindIII-PrA-R (5’-cctgat<u>aagcttg</u>aattcaggatcgtctttaaggcttt-3’) and swapped with GFP into HindIII site of pFRT-TO-DEST-GFP to generate pFRT-TO-DEST-PrA. pENTR4-NOL12 and pFRT-TO-DEST-PrA were recombined with LR clonase to generate pFRT-TO-PrA-NOL12 according to the Gateway system instructions (Life Technologies). The same way, pFRT-TO-Flag-HA-NOL12 was generated by recombining pENTR4-NOL12 and pFRT-TO-DEST-Flag-HA, and pFRT-TO-PrA by recombining pENTR4 with pFRT-TO-DEST-PrA.

### Recombinant protein expression and purification

BL21 (DE3) pLysS cells transformed with pET21a-NOL12-6xHis were grown at 37°C up to OD_600_= 0.6. Cells were induced with IPTG (0.5 mM) for 1 h and harvested by centrifugation. The pellet was flash frozen in liquid N_2_ and stored at ‐80°C. The frozen pellet was thawed and lysed by vortexing in seven pellet volumes of (50mM Tris pH 8, 150 mM NaCl, 10 mM imidazole, 8M urea) until homogenous. The lysate was further homogenized using a Polytron homogenizer (Kinematica PT 1200 E) equipped with a 7mm probe at ¾ maximum speed. The lysate was passed 10× through a 18G ½ needle connected to a syringe. The cell lysate was cleared by centrifugation at 16,000 g for 20’ at 4°C and filtered with a 0.45 μm syringe filter. The cleared lysate was loaded in a 50 mL superloop connected to an AKTA purifier 10 FPLC system (GE Healthcare) and loaded on a 1 mL His-Trap HP column at a flow rate of 1 mL/min. The columns were washed with 10 mL of lysis/binding buffer, and the resin-bound rNol12-His_6_ was refolded using a 20 mL gradient to 100 % (50 mM Tris pH 8, 150 mM NaCl, 10 mM imidazole). rNol12-His_6_ was eluted with 5 mL of (50 mM Tris pH 8, 150 mM NaCl, 350 mM imidazole). Imidazole was removed with a Hiprep 26/10 column equilibrated in (20 mM Tris pH 7.6, 150 mM NaCl, 5 mM MgCl_2_) at a flow rate of 5 mL/min. The sample was concentrated with an Amicon Ultra-15 (3000 NMWL), protein concentration was measured using the Bradford method, and the samples were analyzed by SDS-PAGE.

### Radiolabeled RNA and DNA

Radiolabeled RNA corresponding to T7 polymerase transcribed pBlueScript KS+ linearized by XbaI was used in the nuclease assay below. Briefly, 1.5 μg of linearized and purified template was transcribed in 20 μl of (40 mM Tris pH 7.9, 7.5 mM MgCl_2_, 10 mM DTT, 10 mM NaCl, 2 mM spermidine, 0.5 mM ATP, 0.5 mM CTP, 5 mM GTP, 0.06 mM UTP, 20 U RNasin (Promega), 20 μ& of [αP^32^]UTP (3000 Ci/mmol; Perkin-Elmer)). Reactions were stopped by adding one-tenth volume of 0.5 M EDTA pH 8. Full length RNAs (35 nt) were purified on 12 % denaturing acrylamide gels and eluted in 400 μL of (0.5 M NH_4_OAc, 10 mM Mg(OAc)_2_, 1 mM EDTA, 0.1 % SDS) overnight. Eluted RNA were further extracted with phenol:chloroform (25:1) and precipitated with ethanol (ice-cold, 100 %). RNA pellets were resuspended in 40 μL H_2_O and counted with a beta counter. For the nuclease assays on ssDNA templates, an oligonucleotide corresponding to transcribed RNA above (5'-gggcgaattggagctccaccgcggtggcggccgct-3') was labeled with [γP^32^]ATP (3000 Ci/mmol; Perkin-Elmer). For labeled dsDNA, a reverse complementary oligonucleotide was hybridized to the labeled oligonucleotide above. The dsDNA was then purified on denaturing acrylamide gels and eluted as described above.

The circular dsDNA template for the endonuclease assay was prepared as follows. An unrelated 572 bp fragment was amplified and radiolabeled with [αP^32^]dCTP (3000Ci/mmol; Perkin-Elmer) by PCR. The free nucleotides and oligonucleotides were eliminated with a PCR cleanup column (Qiagen) and the labeled amplicon eluted in 49μl. The cleaned up labeled DNA was phosphorylated in 55μl with 10U of T4 PNK in 1X T4 DNA ligase buffer at 37°C for 30 minutes, and subsequently self-ligated in 300μl of 1X T4 DNA ligase buffer with 6000U of T4 DNA ligase at 16°C for 16 hours. The labeled DNA mixture was separated on a 4% denaturing acrylamide gel, and the circular dsDNA band was excised and eluted, precipitated and quantified as described above.

Circular RNA was prepared from pBS-U17 (26) digested with EcoRI and translated with T7 RNA polymerase (NEB) using 100μCi of [αP^32^]CTP (3000 Ci/mmol; Perkin-Elmer), 16.6μΜ CTP, 0.5mM UTP, 0.5mM ATP, 0.15mM GTP and 0.75mM GMP. Full length RNA was gel purified and circularized by gap-splint ligation with the following oligonucleotide; 5’-ggagacaaaccatgcaggaaacagttttcgaattgggtaccgggccccccc-3’ (27, 28). Equal molar amount of RNA and oligonucleotide (1pmol of each in a volume of 100μl) were denatured for 1 minute at 90°C and cooled to 22°C over a period of 15 minutes. T4 RNA ligase buffer was added to 1x, DMSO to 10%, ATP to 1mM, RNasine (promega) to 1U/μl and T4 RNA ligase to 1U/μl in a final volume of 175μl. The mixture was cooled from 22°C to 16°C over a period of 3 hours, and incubated at 16°C for 16 hours. The circular RNA was then gel purified as described above.

### Filter binding assay

RNPs were reconstituted for 30 minutes at room temperature using 0.2μΜ rNol12-His, 0.3μΜ BSA and 10nM Cy5.5-RNA (5’-Cy5.5gagcucuuccgcagu-3’) in 20μl of 20mM Tris pH7.6, 50mM KCl without MgCl_2_. A negative control was performed in the absence of rNol12-His, and a 5-fold excess of tRNA (50nM) compared to the Cy5.5-RNA was used as a non-specific competitor for rNol12-His binding. During the incubation time, a nitrocellulose membrane was pre-equilibrated in the same buffer for 20 minutes, and placed in a Bio-Dot apparatus (BioRad). The samples were then filtered through the nitrocellulose membrane, and retained samples on the membrane were washed with 40ul reconstitution buffer. The retained RNA on membrane was then resolved on a Li-Cor Odyssey Infrared scanner. The membrane was then blocked in 5% milk and blotted with an anti-His antibody (ABM; G020) to quantify the amount of Nol12 retained on the membrane. RNA corresponding to 1 ul of each reconstitution input were spotted on a nylon membrane as a loading control and scanned using a Li-Cor Odyssey. Signals for retained RNAs were normalized on the signal observed for the BSA only control. The difference in the amount of rNol12-His retained on the membrane was then taken into account to adjust the normalized values for samples with and without tRNA.

### Nuclease assays

Nuclease assay reactions were carried out in 20μl total volume of (20 mM Tris pH 7.6, 150 mM NaCl, 5 mM MgCl_2_). The concentration of rNOL12 either varied (0, 0.1, 0.375, 0.75 and 1.5 μM) in reactions incubated at 30°C for 30 minutes, or 1.5 μM was used for different times (0’, 2’, 10’, 30’) at 30°C. In the divalent cation assay, 5 mM MnCl_2_ or 50 mM EDTA were also used instead of MgCl_2_. In all cases, 10nM of internally radiolabeled RNA, labeled deoxyoligonucleotides or Cy5.5-labeled RNAs were used for the assays. In the case of circular labeled dsDNA, 0.μ1M of DNA was used in the assay. At the end of each reaction, 1 μl of each reaction mixture was added to 9 μl of RNA loading dye (95 % formamide, 0.025 % bromophenol blue, 0.025 % xylene cyanol, 0.025 % SDS, 0.5 mM EDTA). Samples were separated on a 12% denaturing poly-acrylamide gel at 250 V at room temperature and directly exposed using a phosphor screen overnight. The phosphor screen was developed on a Storm 860 Imager.

### Cell Culture and Drug Treatment

HCT116 WT, HCT116 *p53*^-/-^, HEK293T and IMR-90 cells were grown in Dulbecco’s Modified Eagle Medium (DMEM) supplemented with 10 % Foetal Calf Serum (FCS) and Penicillin/Streptomycin (1000 IU, 1 mg/mL respectively; Wisent) at 37 °C with 5 % CO2.

To create the PrA-NOL12 Flp-in cell line, pFRT-TO-PrA-NOL12 and pOG44 expressing the Flp recombinase were co-transfected in HEK293T Flp-In^TM^ cells containing a single FRT recombination site according to the Flp-In^TM^ T-RE×^TM^ core kit manufacturer’s instructions (Life Technologies). Successful transformants were identified by a combination of hygromycin resistance and zeocin sensitivity and doxycycline-inducible expression of PrA‐ or PrA-Nol12 was verified by western blotting. PrA and PrA‐Nol12 expression was induced in the relevant cell lines with 1 μg/mL doxycycline for 24 h and monitored by western blotting using anti-PrA antibodies.

Synthetic siRNAs against NOL12, XRN2, p21, p27 and p53 and a scrambled (SCR) control were supplied by Sigma-Aldrich. siRNAs were transfected using Lipofectamine RNAiMAX (Life Technologies) in a reverse transfection protocol according to the manufacturer’s instructions. Between 7.5×10^4^ and 5.4×10^5^ cells/mL were transfected, depending on the duration of knockdown. Successful silencing of siRNA targets was monitored by western blotting. All siRNA sequences are listed in Supplementary Table 7. Caffeine (2 or 5 mM), VE822 (1 μM), KU55933 (1 μM), ChiR-124 (100 nM), NSC109555 (5μM) or NU7441 (0.5 μM) were added to the above growth media for the duration of the time course; these supplemented media were changed every 24 h throughout the time course to maintain an effective dose. Actinomycin D (0.05 μg/mL or 1 μg/mL) was applied to cells for 3h immediately prior to downstream processing.

### Western Blotting

siRNA-transfected cells were harvested at the indicated time points by trypsin-treatment, washed with PBS and lysed by sonication (5 min 30” on/off, 4°C) in RIPA buffer (50 mM Tris, 150 mM NaCl, 0.1% SDS, 0.5 % sodium deoxycholate, 1 % Nonidet P-40, pH 7.5) supplemented with cOmplete protease inhibitor, EDTA-free (Roche). Soluble fractions were resolved on a 4-12% gradient Bis-Tris NuPAGE^®^ gel according to the manufacturer’s instructions, transferred to a PVDF membrane which was blocked with 5 % milk or BSA in TBS+T (50 mM Tris, 150 mM NaCl, 0.05% Tween-20, pH 7.6) and probed with the indicated primary and secondary antibodies according to Supplementary Table 6. Secondary antibody-conjugated fluorescence was detected using the Li-Cor Odyssey Infrared scanner.

### ssAP-MS/MS analysis of Nol12-containing complexes

HEK293T Flp-In T-REx PrA-Nol12 cells were grown to ~50% confluency, induced for 24 h with 1 μg/mL doxycycline and PrA-Nol12-containing complexes were lysed by cryogrinding (29). PrA-Nol12-containing mRNPs and those from negative controls (PrA-containing or non-epitope tagged) purifications were performed in duplicate with five volumes of extraction buffer (20 mM HEPES-KOH pH 7.4, 0.5% Triton, 1× mini complete EDTA-free protease inhibitor cocktail, 1:5000 antifoam A) supplemented with 100 mM NaCl (+/-100 μg/mL RNAse A) or 300 mM NaCl (+100 μg/mL RNAse A) as previously described (29). Briefly, 0.2 g of cryo-lysed cell powder was slightly thawed (1 minute on ice) and vortex mixed in extraction buffer. The samples were then sonicated (20 W, 2 minutes, on ice). RNAse-treated samples were incubated for 10min at room temperature. All samples were then centrifuged (10 minutes, 4 °C, 16000 × g) and the supernatant was then incubated (30 minutes, slow rotation, 4°C) with 7.5 mg of Dynabeads conjugated with rabbit IgG (160 μg IgG/mg beads) previously washed with extraction buffer. Prior to on-bead digestion, the beads were washed ten times with extraction buffer; once with 100 mM NH4OAC/0.1 mM MgCl^2^/0.5% tritonX-100 (RT, slow rotation, 5 minutes); four times with 100 mM NH_4_OAC /0.1 mM MgCl_2_ (three times fast and once at RT, slow rotation, 5 minutes); and once with 20 mM Tris-HCl pH8.0. Subsequent to ssAP purification, one tenth of each affinity purification sample was eluted with 0.5 M NH_4_OH and resolved on 4-12% gradient Bis-Tris NuPAGE^®^ gels for silver staining. The remaining samples were on-bead trypsin digested in a volume of 50μl (20 μg/mL trypsin in 20 mM Tris-HCl pH 8.0, 37 °C, 900 rpm, 16-20 h; stopped with 2 % formic acid; (30)) and analyzed by tandem mass spectrometry as described previously (31).

Nol12-interacting proteins were identified as follows. The highest total spectrum count values of the controls (Untagged, PrA; Supplementary Table 1) were subtracted from that of the PrA-Nol12 ssAP samples under the respective condition. A peptide was considered enriched in a PrA-Nol12 ssAP sample (100/300mM NaCl +/-RNAse A) if the spectrum count value for each replicate was ≥2-fold greater than the highest spectrum count value recorded among the relevant controls (PrA and untagged) under the same purification conditions. The total number of unique peptide spectra recorded for each prey was then determined; preys were defined as Nol12-associated proteins if at least three enriched peptides were identified in both experimental duplicates for one or more of the purification conditions (+/-RNAse A, +/-supplemental NaCl). For each prey protein identified, the number of unique peptide spectra recorded was normalized against the number of bait protein peptides recorded; this relative number of unique peptide spectra recorded per protein was used as a metric for semi-quantitative analysis of Nol12-associated proteins.

### MS analysis of rNol12

A total of 1ug of purified recombinant Nol12 was denatured in 6M urea, 100mM ammonium bicarbonate at 37°C for 10 minutes, then reduced with 10mM DTT at 37°C for 30minutes before being alkylated with 16.6mM IAA for 1h in the dark at room temperature. Urea was then diluted to 1.6M and ammonium bicarbonate adjusted to 75mM. Trypsin was added in a 1:20 trypsin:protein ratio and incubated overnight at 37°C. Next day, the sample was dried using a speedvac, resuspended and cleaned with a ZipTip C_18_ according to the manufacturer (EMD Millipore) before being analyzed by tandem mass spectrometry as described previously (31).

### Northern blotting

Total RNA from scramble and NOL12 siRNA transfected cells was extracted with TRIzol (Invitrogen) according to the manufacturer’s protocol. For long RNA analysis, total RNA were separated on 1% agarose-formaldehyde gels in Tricine-Triethanolamine and transferred on nylon membrane in 10x SSC by capillarity as described by Mansour and Pestov (32). Small RNAs were separated on 8% acrylamide-urea gels and transferred on nylon membrane in 0.5× TBE overnight at 12 V. For long rRNA analysis, 3 μg of total RNA extracted from HCT116 WT and HCT116 p53^-/-^ cells were used, as well as 6 μg of RNA extracted from IMR-90 cells. A total of 8 μg of RNA was used for small RNA analysis. All probes used for hybridization are shown in Supplementary Table 8. Probes identified with terminal amines were fluorescently labeled with DyLight^TM^ 800 NHS Ester as previously described (33), while other probes were radiolabeled with [γP^32^]ATP (3000 Ci/mmol; Perkin-Elmer).

### Cell cycle profiling and quantification of apoptosis

To profile cell cycle distribution, siRNA‐ and/or drug-treated cells were harvested by trypsinization at the indicated time points, washed twice with PBS and fixed with 70% ethanol at ‐20 °C for ≥1 h. Fixed cells were rehydrated 2× in PBS then permeabilized, RNAse-treated and stained for DNA content with a modified Krishan Buffer (20 μg/mL propidium iodide, 0.1 % Triton X-100, 0.2 mg/mL RNAse A in PBS) for 30 min at RT. Apoptosis was measured using the Annexin V-FITC apoptosis detection kit (Sigma-Aldrich) according to the manufacturer’s instructions. Briefly, siRNA‐ and/or drug-treated cells were harvested at the indicated time points and resuspended at 5x10^5^ cells/ml. Cells were treated with Annexin V-FITC (250 ng/ml) and propidium iodide (500 ng/ml). In both cases, staining was quantified by flow cytometry and results were analyzed using the FlowJo software.

### Detection of oxidized DNA

siRNA-transfected cells were harvested at the indicated time points and DNA was extracted as previously reported (34). A total 2 μg of DNA per sample was spotted onto a nylon membrane and stained with methylene blue, before being digitally scanned to serve as loading control. The membrane was then blocked overnight with 1% casein in SSC buffer. To detect oxidized guanosines, the membrane was incubated overnight at 4°C with an anti-8-OHdG antibody (Millipore). The membrane was then washed 3 times in TBST and incubated with a IRDye 800CW fluorescently labeled donkey anti-goat secondary antibody diluted 1:15000 in TBST for 1 h. The membrane was finally scanned with a Li-Cor Odyssey infrared scanner. As positive control, a Fenton reaction consisting of 0.5 μg/μl genomic DNA was incubated with 25 μM CuSO_4_ and 50 mM H_2_O_2_ for 1 h at 37°C.

### Immunofluorescence

siRNA-transfected or doxycycline-induced cells were grown on poly-L-lysine-coated glass coverslips for 48 h then fixed with 3.2 % paraformaldehyde (10 min). Cells were washed 2× with PBS, permeabilized with 0.25% Triton X-100 in PBS (15 min) and blocked with 1 % BSA in PBS+T (50 mM Tris, 150 mM NaCl, 0.05% Tween-20, pH 7.6). The indicated primary antibodies were hybridized for 1 h in 1% BSA in PBS+T, cells were washed 3× with PBS then rehybridized with the relevant secondary antibody in the same conditions. Coverslips were washed a further 3× with PBS, DNA was stained with Vectashield mounting medium containing DAPI (Vector Laboratories) and imaged using the Zeiss Axioimager Z2 upright epifluorescence microscope with a 63× objective, fitted with the Zeiss Axiocam mRm CCD camera. Images were processed with FIJI open-source image analysis software (35). Traces were generated with the ‘Plot Profile’ function in Fiji using representative images from each immunofluorescence experiment. Profiles for each channel were normalized against the maximum observed fluorescence value along the line segment and colocalization determined using the Fiji software Coloc2 module.

## RESULTS

### Nol12 associates with a diverse interactome

In order to more thoroughly investigate Nol12’s cellular niches, we determined its interactome using Protein A-tagged Nol12 (PrA-Nol12) expressed in HEK293T cells by single-step affinity purification (ssAP) coupled to semi-quantitative mass spectrometry (MS), and analyzed it against untagged and PrA-expressing cell lines (Supplementary Figure 1a) (31, 36). The highest total spectrum count values found in the control samples (Untagged, PrA; Supplementary Table 1) were subtracted from that of a PrA-Nol12 sample under the respective condition (100 or 300mM NaCl +/-RNAse A). A peptide was considered ‘real’ in a particular PrA-Nol12 sample (100 or 300mM NaCl +/-RNAse A) if the spectrum count value in each replicate was ≥2-fold greater than the highest spectrum count value recorded among the relevant controls under the same purification conditions (Supplementary Tables 1 and 2). The total number of unique peptide spectra recorded for the remaining preys was determined, and preys were defined as Nol12-associated proteins if at least three enriched peptides were identified in both replicates for each condition. For each prey protein identified, the number of unique peptide spectra recorded was normalized against the bait protein peptide count to obtain a number of unique peptide spectra per prey, which was used as a metric for analysis of Nol12-associated proteins. We identified a total of 759 proteins, of which 496 were defined as enriched (≥3 peptides in all replicates; Supplementary Table 2). Functional classification of the identified proteins showed a complex interactome and suggested a highly diverse functionality for Nol12 in human cells (Supplementary Tables 3 and 4).

In line with a putative nucleolar and ribosome biogenesis-associated role, PrA-Nol12-associated preys included 75 ribosome maturation factors required for large ribosomal subunit (LSU) maturation and 54 known 90S pre-ribosomes components, while only 8 proteins specific for small ribosomal subunit (SSU) maturation were identified (Figure 1a). In addition, 42 LSU and 27 SSU ribosomal proteins (RPs) were found (Figure 1a), suggesting a role for Nol12 during LSU maturation events. Comparison of our set of Nol12 interactors with human ribosome biogenesis proteins identified in a recent high-throughput study showed a 58.9% (n=158) overlap supporting the idea of Nol12 as a ribosome maturation factor (Supplementary Figure 1b) (37). However, PrA-Nol12 also co-isolated a significant number of proteins from several non-ribosomal processes. Those included 51 proteins involved in genome maintenance and integrity (Figure 1a, b: ‘Chromatin associated’; Supplementary Table 3), including Dh×9, PRKDC and MYBBP1A. Furthermore, 94 mRNA maturation and turnover factors were identified, including splicing and mRNA maturation factors (e.g. snRNP200, Srsf1, Thoc2, Mrto4), paraspeckle proteins (NONO, SfpQ) as well as components of processing (P) bodies, sites of post-transcriptional mRNA regulation in the cytoplasm (e.g. Stau1, hnRnpA3, Fmr1, LaRP1) (Figure 1a, b: ‘mRNA metabolism’; Supplementary Table 3) (1). Moreover, we identified 69 proteins with functions in mitochondria, including many involved in mitochondrial ribosome biogenesis (Figure 1a, b: ‘Mitochondria’; Supplementary Table 3). The observation that MRPLs/MRPSs represent low-frequency contaminants in affinity purification experiments argues against these interactions arising from non-specific interactions (Supplementary table 1) (38).

**Figure 1.**
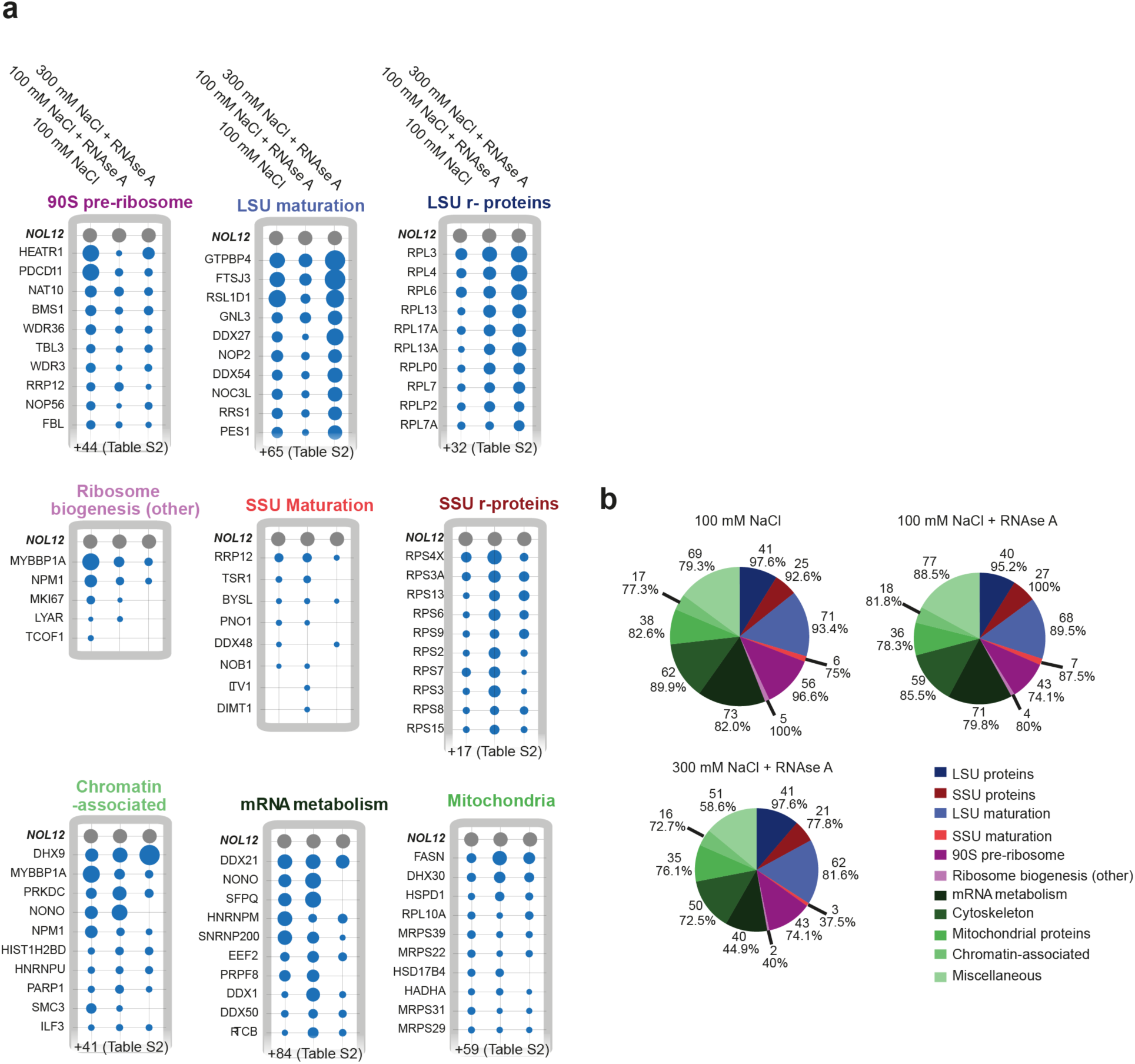
Nol12 associates with a diverse range of complexes. PrA-Nol12-associated complexes were affinity purified from HEK293T cells. Peptides enriched more than 2-fold over negative controls (untagged and PrA-only cell lines) and present in ≥3 peptides in all replicates, were identified and mapped to the human proteome. (a) Total unique peptide counts for enriched peptides were normalized against bait peptide counts, and resulting relative peptide counts for each condition were plotted as radii of a circle (10 most-enriched proteins shown here). (b) The numbers of proteins from each functional class (based on GO terms) found associated with PrA-Nol12 under each purification condition were calculated. Percentages represent the relative coverage of proteins from the particular functional class identified under that condition; pie graph sizes are proportional to the total size of the PrA-Nol12 interactome under that condition.

To determine the dependency of Nol12 associations on RNA as well as their stability, we furthermore investigated the Nol12 interactome in the presence of RNase A and/or increased salt. Components of the early (90S) pre-ribosomes were reduced by RNAse digestion, while the majority of LSU maturation factors and RPLs were considerably less sensitive to RNAse treatment (Figure 1a, b; Supplementary Table 3), suggesting an RNA-dependency of Nol12 recruitment to 90S pre-ribosomes over 60S pre-ribosomes; however, an incomplete RNA digest due to the protection of pre-rRNA within LSU pre-ribosomal complexes cannot be ruled out. LSU maturation factors and RPLs were also considerably stabilized under increased salt conditions (Figure 1b). Of the non-ribosome complexes, only a small number of co-isolated ‘RNA metabolism’ and ‘chromatin-associated’ factors showed sensitivity to RNAse treatment, suggesting that most Nol12 interactions in this group are not RNA-dependent (Figure 1b); however, the majority of ‘mRNA metabolism’ interactors exhibited salt sensitivity compared to the ‘chromatin-associated’ group, suggesting labile or transient associations of Nol12 with interactors involved in mRNA metabolism.

### Nol12 localizes to different subcellular compartments

Proteomic analysis of Nol12-associated complexes identified both nuclear and cytoplasmic proteins involved in nucleic acid homeostasis, supporting previous high through-put data sets on Nol12 localization (23). To more concretely define these subcellular compartments, we carried out fluorescent microscopy in HCT WT cells using an anti-Nol12 antibody, and compared the protein’s localization to that of established markers for different subcellular structures. We found a large fraction of Nol12 localized to nucleoli, with a small amount in the dense fibrillar component (DFC), marked by co-staining for the early ribosomal biogenesis factor Fibrillarin (39), while more Nol12 was found within the granular component (GC), consistent with an 90S pre-ribosomes association and a role in early LSU maturation (Figure 2a). In addition to its nucleolar localization, Nol12 was also present within numerous discrete foci throughout the nucleoplasm (Figure 2a), some of which showed distinct overlap with the paraspeckle component SfpQ (Figure 2b), concurrent with our AP-MS data (Figure 1a), and consistent with previously observed SfpQ localization patterns (40, 41). Further co-staining of Nol12 with markers for different nuclear structures indicated that the protein was most likely not a component of polycomb complexes (Bmi1; Supplementary Figure 2b), Cajal bodies (Coilin; Supplementary Figure 2c), or PML bodies (PML; Supplementary Figure 2d) as no spatial overlap with respective markers was determined using Fiji’s Coloc2 module analysis (Materials and Methods). Some partial overlap was found with slicing speckles, (SC35; Supplementary Figure 2a), which could be due to Nol12’s localization to paraspeckles, which are found adjacent to splicing speckles (42).

**Figure 2:**
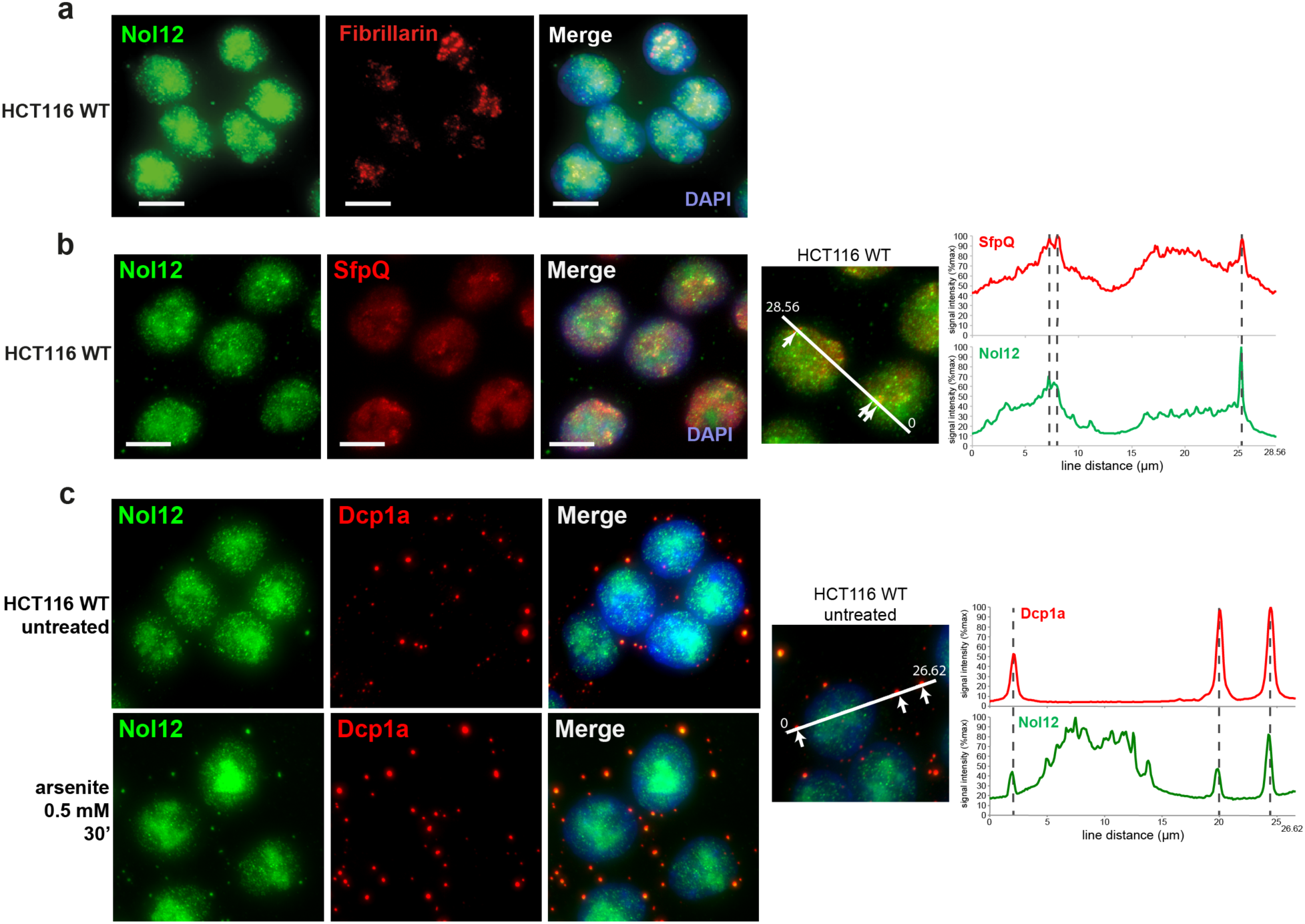
Nol12 co-localizes with nucleoli, paraspeckles, mitochondria and GW/P-bodies. Endogenous Nol12 localization in HCT116 WT cells was compared to that of the nucleolar marker Fibrillarin (a), the paraspeckle component SfpQ (b), and the GW/P-body component Dcp1a (c). Images are representative of three independent experiments. Scale bar = 10 μm. For (c) HCT116 WT cells were grown on poly-L-lysine-coated coverslips for 48h, with or without treatment with sodium arsenite (0.5 mM, 30’) prior to fixation, imaging and trace analysis. (b, c) Linear profiles of fluorescence density were created from representative images of each co-immunofluorescence experiment and normalized against the maximum observed fluorescence along the line segment. Arrows/dotted lines indicate the location of local fluorescence maxima in the marker channel; colocalization was analysed using the Fiji software Coloc2 module.

Nol12-associated complexes identified by MS also included cytoplasmic components, in particular GW/P-body proteins (Figure 1a, b). Using an anti-Nol12 antibody, we observed several bright, discrete cytoplasmic foci in HCT116 WT cells (Figure 2a, c), which overlapped with co-staining for Dcp1a, a key component of the mRNA-containing GW/P-bodies, and were significantly increased after treatment of cells with arsenite (Figure 2c) (43). Interestingly, under arsenite conditions, Nol12 was not found localized to cytoplasmic stress granules (Tia1; Supplementary Figure 2e). Localization of Nol12 to these various substructures was observed using different anti-Nol12 antibodies, raised against different portions of the protein (Sigma aa130-215; Bethyl aa75-115; data not shown). Taken togther, the localization of Nol12 to different subcellular compartments not only supports our proteomic data, but also suggests multiple functions for the protein.

### Nol12 exhibits RNA endonuclease activity *in vitro*

The S. *cerevisae* homologue of Nol12, Rrp17, has previously been shown to possess RNA binding and 5’-3’ RNA exonuclease activity (22). To determine whether Nol12 can bind and process RNA, and to better characterize Nol12’s role within its diverse interactome, we tested its nuclease activity *in vitro* using different RNA substrates. Purified recombinant Nol12 (rNol12) was able to degrade a short, uniformly labeled RNA substrate into a ladder of truncated products in a concentration-dependent manner *in vitro* (Figure 3a). In line with its nuclease activity and high proportion of positively charged residues, Nol12 exhibited RNA binding activity, however, RNA binding was not specific as demonstrated by competition with tRNA (Supplementary Figure 3b). Next, we asked if Nol12 had a specific requirement for divalent cations. Using short, uniformly labeled RNA substrates, the nucleolytic activity of Nol12 was inhibited by replacement of MgCl_2_ with MnCl_2_ or pretreatment with EDTA (Figure 3b). To examine direction-dependent degradation, we used an *in vitro* transcribed single-strand (ss) RNA labeled at its 5’ or 3’ end. Incubation of rNol12 with 5’ end-labeled RNA resulted in a gradual loss of the full-length labeled substrate, however, no accumulation of radiolabeled nucleotide was detected; instead the appearance of radiolabeled products of different lengths was observed (Figure 3c, left). Similar observations were made upon incubation of rNol12 with 3’ end-labeled substrate (Figure 3c, right). This suggests that the RNA products are the result of endonucleolytic cleavages rather than end-degradation, which led us to test Nol12 for *in vitro* endonucleolytic function. Upon incubation of circular radiolabeled ssRNA with rNol12, the circular RNA was linearized producing both a linear ssRNA followed by steady degradation over the course of 30 minutes, resulting in a smear of smaller products (Figure 3d); the appearance of the latter could either be due to unspecific endonucleolytic cleavage, or and additional exonucleolytic activity, which cannot be excluded. Taken together, these results suggest that Nol12 has endonucleolytic activity *in vitro*. However, rNol12 did display low efficiency in vitro; after the exclusion of contaminating nucleases co-purified with rNol12 by mass spectrometry and PAGE analysis (Supplementary Figure 3a; Supplementary Table 5), it suggests that Nol12 may require a co-factor for efficient nuclease function. Finally, given the exonucleolytic activity of the Nol12 yeast homologue Rrp17, a similar activity for Nol12 cannot be complete ruled out; conversely, Rrp17 has never been tested for endonuclease function.

**Figure 3.**
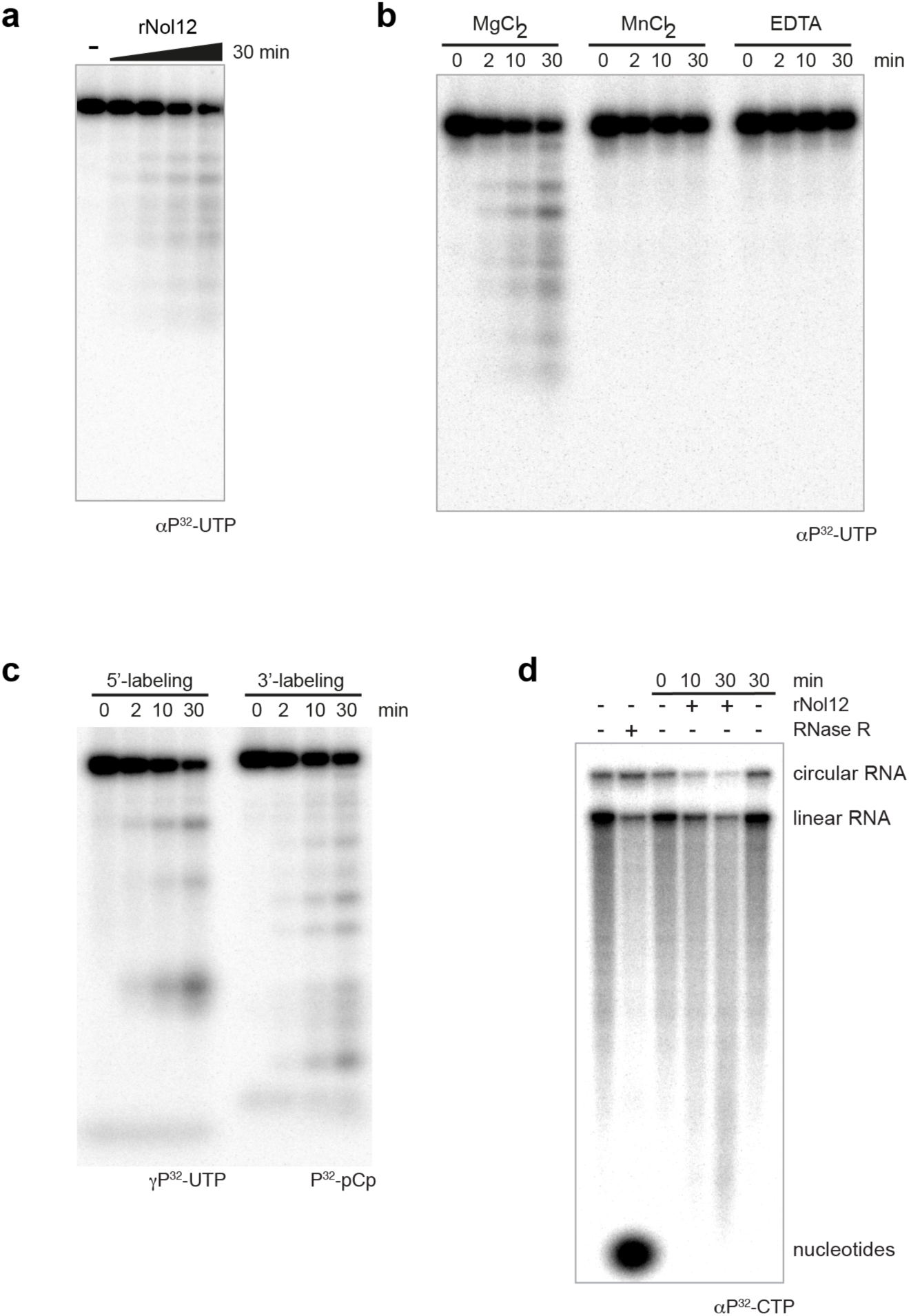
Nol12 is an RNA endonuclease. (a) Nuclease assays using *in vitro* transcribed radiolabeled RNA (αP^32^-UTP) and recombinant rNol12. Increasing rNol12 concentrations up to 1αM were incubated at 30 °C for 30 minutes before being resolved on a 12% denaturing acrylamide gel. (b) Recombinant Nol12 was incubated with *in vitro* transcribed radiolabeled RNA (αP^32^-UTP) at 30 °C for 0, 2, 10 and 30 minutes in the presence of MgCl_2_, MnCl_2_ or excess EDTA before being resolved on a 12% denaturing acrylamide gel. Images are representative of three independent experiments. (c) Degradation of *in vitro* transcribed 5’-γP^32^-ATP (left) or 3’-pCp ‐αP^32^ (right) end-labeled ssRNA by rNol12. (d) Nuclease assay performed on a mixture of linear and circular RNA. RNA alone (lane 1) was treated with RNase R (lane 2) exonuclease. The RNA was incubated in nuclease buffer without rNol12 for 0 and 30 min (lanes 3, 6), and with rNol12 for 10 and 30 minutes (lanes 4-5).

### Nol12 is required for rRNA processing at site 2 during ribosome biogenesis

It has previously been suggested, based on studies in HeLa cells, that Nol12’s presence on preribosomes may be required for processing events within the Internal Transcribed Spacer 1 (ITS1) (4). Processing of ITS1 requires three endonucleolytic cleavages; however, only endonucleases for two of these sites, Rcl1 and Nob1, have been identified so far, which act in concert with the exonucleases Xrn2 and Rrp6 (4, 37). Given the *in vitro* endonucleolytic activity of Nol12, we investigated the function of Nol12 in pre-rRNA processing in more detail. HCT116 wild-type (HCT WT) cells were transfected with an siRNA targeting NOL12 transcripts (siNOL12 #5; referred to as siNOL12 throughout unless otherwise stated) (Figure 4a, lanes 2, 4 and 6), or a scrambled control siRNA (siSCR) (Figure 4a, lanes 1, 3 and 5), and pre-rRNA processing was analyzed by Northern blotting 24h, 36h and 48h after transfection. Knockdown of NOL12 in HCT WT cells resulted in rRNA processing defects affecting precursors produced by cleavage at site 2 within ITS1: the large subunit precursor 32.5S, and the small subunit pre-rRNAs 30S, 26S and 21S were noticeable decreased after 36h compared to control (Figure 4a). Conversely, levels of site E cleavage products, namely 36S pre-rRNAs, its intermediate processed form 36S-C as well as the small subunit precursor 18S-E, were increased as early as 24h post siRNA treatment (Figure 4a). Mature rRNAs levels were overall only modestly decreased, with LSU rRNAs more affected than SSU ones, suggesting a rerouting of the ribosome maturation pathway to sustain ribosome production. The same modulation of precursors was also observed in IMR-90 primary fibroblasts (Supplementary Figure 4a, lanes 2 and 4), although with differing kinetics most likely due to the cells’ overall slower growth rate.

**Figure 4.**
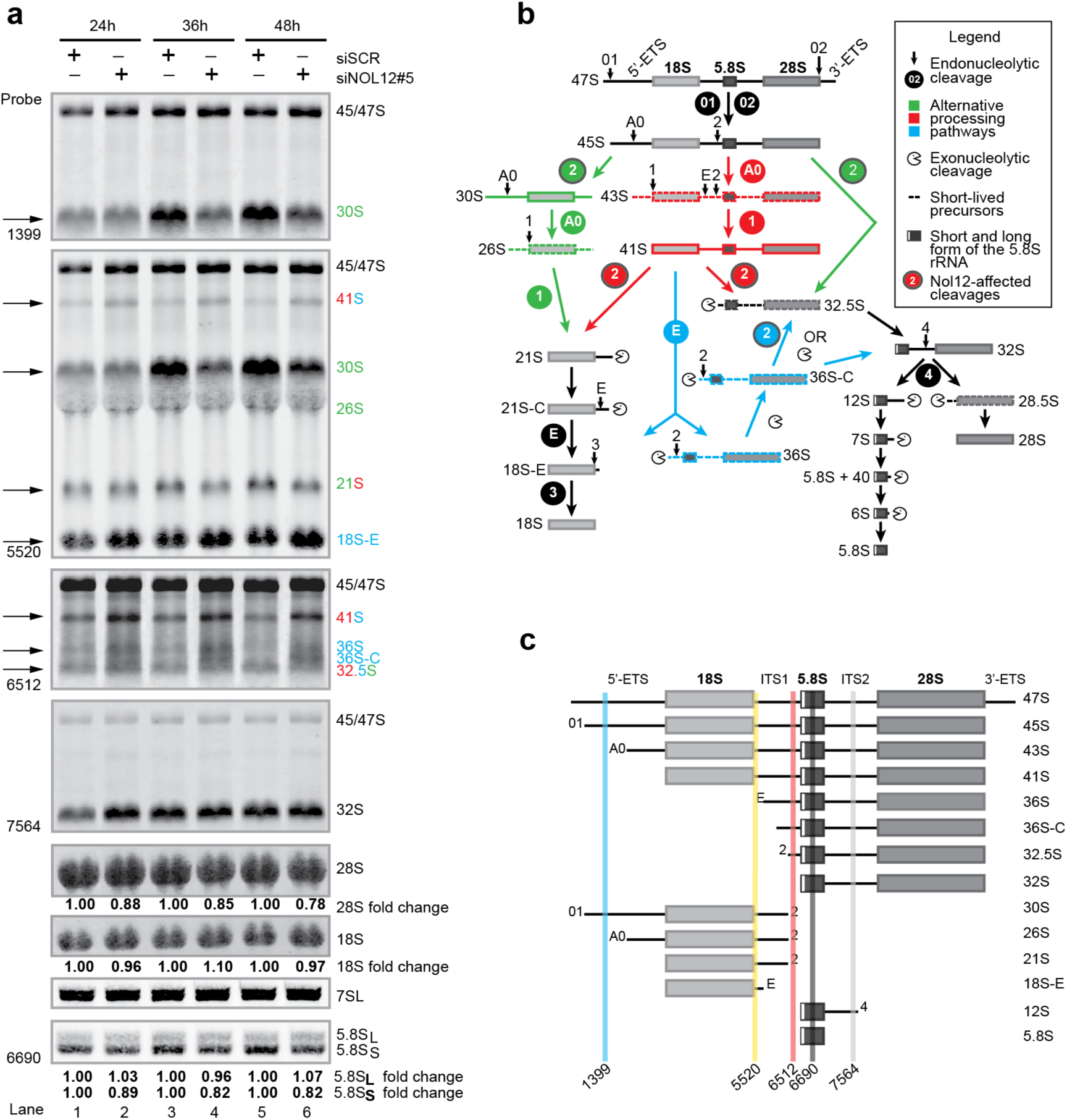
Nol12 is required for large and small ribosomal subunit maturation. (a) Total RNA extracted at different time points from HCT116 WT cells transfected with siNOL12#5 or siSCR was analyzed by Northern blotting using probes hybridizing to either 5’-ETS, ITS1 or ITS2 regions of the pre-rRNA to detect processing intermediates. RNA loading was monitored by methylene blue staining of mature 18S and 28S rRNAs (large RNAs) or Northern blotting for the 7SL RNA (small RNAs). A schematic of rRNA maturation pathways in human cells is shown in (b), and the relative positions of probes used with respect to different rRNA processing intermediates are shown in (c).

In higher eukaryotes, processing of Pol I-transcribed pre-rRNA occurs along three parallel pathways, presumably to ensure biogenesis of ribosomes (Figure 4b). Pathway 1 and 2, the major pathways, require an endonucleolytic cleavage in ITS1 at site 2 to separate SSU and LSU pre-rRNAs (Figure 4b; green and red, respectively), while the alternative, minor pathway 3 produces downstream precursors via endonucleolytic cleavage at site E (Figure 4b; blue) (2). Pre-rRNAs generated upon site 2 cleavage (30S, 32.5S, 21S) are all decreased in the absence of Nol12, while pre-rRNAs that are either substrate (41S) for, or products of site E cleavage along pathway 3 (36S and 18S-E) are increased (Figure 4a, arrows). This increase of 41S and 18S-E, together with a concomitant decrease of 21S (product of site 2 cleavage) and appearance of 36S and 36S-C (products of site E cleavage; Figure 4a), indicates that kd of NOL12 leads to an overall decreased cleavage of ITS1 at site 2, and shift of processing events to site E (Figure 4b, blue), suggesting a requirement for Nol12 for processing at site 2. We did not detect reduced processing at site E or within 5’ETS, events which have been attributed to the endonuclease Rcl1 and the 5′-3′ exonuclease Xrn2, respectively; moreover, the precursors 36S and 36S-C, which are increased upon NOL12 kd, are known substrates for Xrn2 in mouse and HeLa cells (4). The only marginally lower levels of 5.8S, 18S and 28S indicate the continued production of mature rRNAs, presumably along pathway 3, to ensure ribosome biogenesis, suggesting that, in the absence of Nol12, the 36S precursor is processed to 32S by Xrn2 following Rcl1 cleavage at site E, albeit with slower kinetics, explaining the accumulation of 41S and 36S-C, while 32S levels are unchanged; 18S-E, also generated by a cleavage site E, is matured to 18S rRNA by Nob1 (44). Taken together, and in light of its RNA endonuclease activity shown above, these data suggest Nol12 as the endonuclease required for ITS1 cleavage at site 2 (Figure 4a; Supplementary Figure 4a).

### Loss of Nol12 leads to p53-independent cell cycle arrest and apoptosis

As Nol12 associated with proteins outside ribosome biogenesis and the nucleolus, and ribosome biogenesis was maintained in the protein’s absence, we went on to investigate the overall effect of Nol12 depletion on cell proliferation and survival in HCT WT cells to obtain information on its other functional niches. Knockdown of NOL12 resulted in a robust G_1_/S arrest after 36-48h (Figure 5a). Strikingly, depletion of Nol12 also induced a strong apoptotic response within 36-48h of siRNA transfection, as evidenced by both increased annexin V binding without concurrent propidium iodide infiltration (Figure 5b), and by proteolytic activation of caspase 3 (Figure 5c). This rapid onset of apoptosis is highly atypical of rRNA maturation factor depletion (37), and, by comparison, knockdown of the multifunctional 5’-3’ exonuclease and ribosome processing factor XRN2 induced only a subtle G_1_/S delay within 24h of siRNA treatment (Figure 5a), and no apoptotic response (Figure 5c) suggesting that the observed apoptotic was not due to Nol12’s role in ribosome biogenesis. Analysis of HCT WT whole cell lysates by western blotting revealed that depletion of Nol12, but not Xrn2, resulted in progressive accumulation of the key cell cycle and apoptosis-regulatory protein p53, as well as p21^WAF1/CIP1^ and p27^KIP1^ (Figure 5d). These changes coincided with downregulation of S-phase-promoting phosphorylation of Rb and increased phosphorylation of p53 on Ser15, a stabilizing modification by PIKK-family kinases (ATM, ATR and DNA-PK) in response to DNA damage (Figure 5d) (45), suggesting that loss of Nol12 elicits significant cellular stress, activating both cell cycle arrest and apoptosis response pathways, despite its modest effect on ribosome biogenesis.

**Figure 5.**
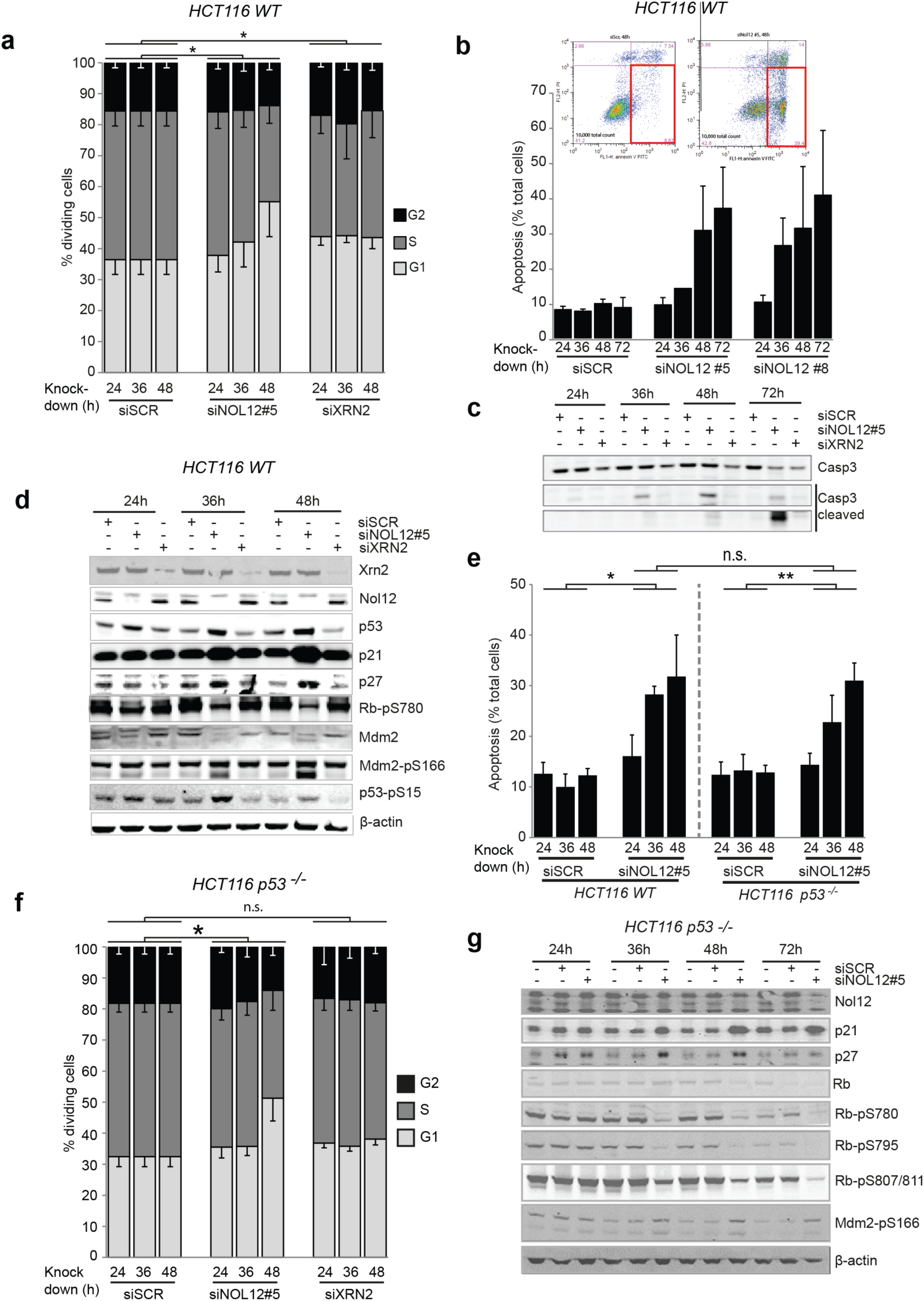
NOL12 knockdown causes p53-independent G_1_/S arrest and apoptosis. (a-d) HCT116 WT cells were transfected with the indicated siRNAs (10μM) for the times specified. (a) Propidium Iodide (PI)-stained DNA content was quantified by FACS, and G_1_ (2n), S (2n-4n) and G_2_ (4n) populations determined. Data represent mean±s.d. for five independent experiments. *p<0.05 by Student’s two-tailed t-test on area under the curve (AUC) of G_1_ or AV+ PI^-^ populations as appropriate. (b) Annexin V-FITC-positive, PI-negative apoptotic cells (gated, inset) were quantified by FACS. Data represents mean±s.d. for two independent experiments. (c, d) Whole cell extracts (WCEs) of siRNA-treated cells were resolved on 4-12% Bis-Tris gradient gels, transferred to PVDF membrane and blotted with the indicated primary antibodies. (e-g) HCT116 *p53*^-/-^ cells were treated with the indicated siRNAs (10μM) for the specified times. (e) HCT 116 WT or *p53*^-/-^ cells were treated with siSCR or siNOL12#5 (10μM) for the times indicated, and apoptotic cells quantified. Annexin V-FITC-positive, PI-negative apoptotic cells (gated, inset) were quantified by FACS. Data represents mean±s.d. for two independent experiments. * p<0.05, ** p<0.05, n.s., non-significant by Student’s two-tailed t-test on area under the curve (AUC) of G_1_ or AV+ PI^-^ populations as appropriate. (f) Propidium Iodide (PI)-stained DNA content was quantified by FACS, and G_1_ (2n), S (2n-4n) and G_2_ (4n) populations determined. Data represent mean±s.d. for five independent experiments. (g) HCT116 *p53*^-/-^ cells were treated with the indicated siRNAs (10μM) for the specified times. Whole cell extracts (WCEs) of siRNA-treated cells were resolved on 4-12% Bis-Tris gradient gels, transferred to PVDF membrane and blotted with the indicated primary antibodies.

Recent data has shown that loss of several ribosome biogenesis factors or ribosomal proteins triggers ‘nucleolar stress’ via accumulation of free 5S RNP (5S rRNA, Rpl5 and Rpl11) in the nucleoplasm, leading to inhibition of ribosome biogenesis and downregulation of the p53-ubiquitin E3 ligase Mdm2. This leads to subsequent p53 accumulation and cell cycle arrest, thus providing cells with a surveillance mechanism for monitoring ribosomal integrity (46). Nol12 depletion did not result in accumulation of nuclear Rpl11 (data not shown), suggesting that NOL12 kd-induced cell cycle arrest and apoptosis may not be part of a nucleolar stress response. To determine whether NOL12 kd-mediated G_1_/S arrest and apoptosis are dependent on p53 stabilization, Nol12 was depleted in HCT116 cells lacking functional p53 (HCT116 *p*53^-/-^(47)). Both apoptosis (Figure 5e) and G_1_/S arrest (Figure 5f) were identical to p53^+/+^ cells upon loss of Nol12, while no cell cycle delay was detected upon XRN2 kd (Figure 5f). Pre-rRNA maturation defects upon NOL12 kd in the absence of p53 were identical to those observed in HCT WT cells (Supplementary Figure 4b), consistent with previous reports regarding ribosome biogenesis in p53-deficient cells (37). Similar results were obtained when Nol12 and p53 were codepleted in HCT WT cells (Supplementary Figure 5a), suggesting that the G_1_/S delay and apoptosis observed upon NOL12 kd are not dependent on p53 signaling.

While p53-dependent pathways represent the major effector mechanisms of nucleolar stress, p21^WAF1/CIP1^, p27^KIP1^ and c-myc were previously shown to activate nucleolar stress-dependent cell cycle arrest in a p53-independent manner (46). Indeed in HCT116 *p53*^*-/-*^ cells, p21^WAF1/CIP1^ and p27^KIP1^ were both upregulated following NOL12 kd (Figure 5g), while c-myc, another target of Rpl11, was unaltered in both HCT WT and *p53*^*-/-*^ cells (Supplementary Figure 5b, c) (46). However, co-depletion of Nol12 and either p21^WAF1/CIP1^ or p27^KIP1^ in HCT *p53*^*-/-*^ cells (Supplementary Figure 5d), or of Nol12, p53 and either p21^WAF1/CIP1^ or p27^KIP1^ in HCT WT cells failed to rescue G_1_/S arrest (Supplementary Figure 5a), suggesting that the observed cell cycle arrest, and apoptosis response, upon Nol12 depletion are neither p53 nor c-myc dependent as well as independent of p21 or p27, and thus qualitatively distinct from a nucleolar stress response. Interestingly, while we did observe downregulation of Mdm2 after loss of Nol12, increased amounts of the phosphorylated form Mdm2^pS166^ were detected in NOL12 kd (Figure 5d), and more mildly in HCT116 *p53*^*-/-*^ cells (Figure 5g), but not XRN2 kd cells (Figure 5d). Mdm2^pS166^ is an activating phosphorylation mediated by p-Akt upon oxidative stress that propagates p53 degradation; however, binding of Mdm2^pS166^ to p53 is blocked by p53-phosphorylation on Ser15/Ser20. Upon increased oxidative stress and Mdm2^pS166^ phosphorylation, p53^pS15^ is degraded, an effect also observed upon NOl12 kd in HCT WT cells after 48h (Figure 5d), suggesting that activation of p53^pS15^ and activation of Mdm2^pS166^ are caused by different upstream events and effectors, and may activate different downstream pathways (45, 48).

### Nol12-induced apoptosis is dependent on the DNA damage-sensing kinase ATR

The induction of a robust, p53-independent apoptosis following Nol12 depletion is highly unusual for a ribosome biogenesis factor, supporting the notion that Nol12 may possess non-ribosomal functions, which are responsible for this apoptotic response. The phosphorylation of p53 on Ser15, an event associated with the activity of the DNA damage-sensing PIKK-family kinases ATM, ATR and DNA-PK, and Mdm2 on Ser166, an event mediated by p-Akt1 upon oxidative stress, following Nol12 depletion, led us to examine the activation of DNA damage response (DDR) pathways. Western analysis of whole cell lysates from NOL12 kd cells revealed the induction of signaling cascades typically associated with a DDR (Figure 6a). Phosphorylation of the variant histone H2A.X on Ser139, an early event in DNA damage recognition, was increased after 24h, an effect also seen upon kd of XRN2 but to a lesser extent. NOL12 kd also resulted in robust and rapid phosphorylation of Chk1 on Ser345 after only 24hr, indicating activation of the ATR kinase (Figure 6a), while only a mild increase in phosphorylation of Chk2 at Thr68 and a decrease of Chk2^pSer19^, targets of DNA-PK and ATM kinases, respectively, were observed, suggesting that loss of Nol12 triggers activation of the ATR-Chk1 arm of the DDR cascade, and, to a lesser extent, ATR-Chk2 (49).

**Figure 6.**
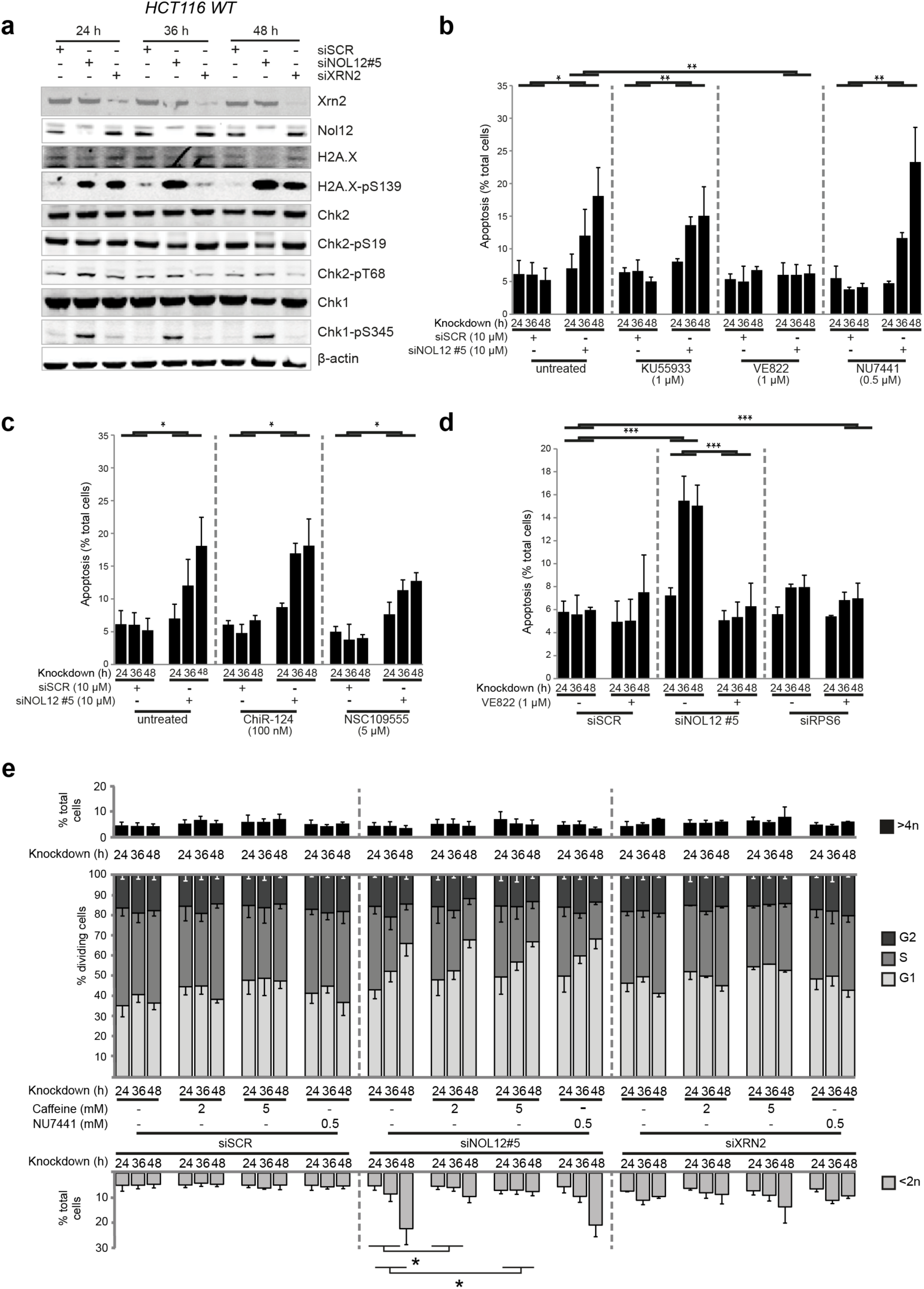
Loss of Nol12 causes apoptosis via ATR signaling. (a) WCEs of HCT116 WT cells treated with indicated siRNAs for the times specified were resolved on 4-12% Bis-Tris gradient gels, transferred to PVDF membrane and blotted with the indicated primary antibodies. (b) HCT116 WT cells were transfected with siSCR or siNOL12#5 (10μM) as indicated and grown in the presence of KU55933 (μM; ATM inhibitor), VE822 (μM; ATR inhibitor) or NU7441 (0μM; PRKDC inhibitor) for the indicated times prior to quantification of apoptotic cells by FACS as in Figure 5b. Data represents mean±s.d. for four independent experiments. (c) HCT116 WT cells were transfected with siSCR or siNOL12#5 (10μM) as indicated and grown in the presence of ChiR-124 (100nM; Chk1 inhibitor) or NSC109555 μM; Chk2 inhibitor) for the indicated times prior to quantification of apoptotic cells as in Figure 4B. Data represents mean±s.d. for four independent experiments. (d) HCT116 WT cells treated with the indicated siRNAs and grown in the presence or absence of VE822 (μM) for the indicated times prior to quantification of apoptotic cells as in (b). Data represents mean±s.d. for four independent experiments. (e) HCT116 WT cells treated with the indicated siRNAs were grown in the presence of caffeine (2 or 5mM) or NU7441 (0μM) for the indicated times prior to analysis as described in Figure 4A except that <2n and >4n populations were also scored. Data represents mean±s.d. for >4 independent experiments. * p<0.05, ** p<0.01, *** p<0.005 by Student’s two-tailed t-test on AUCs of AV+ PI^-^ or <2n populations as appropriate.

ATM, DNA-PK and ATR are activated by different DNA damage signals. ATM and DNA-PK recognize double-strand breaks (DSBs), while ATR recognizes regions of single-strand DNA that arise from a range of insults including single-strand breaks (SSBs), DNA oxidation and stalled replication forks; however, some cross-talk between the pathways exists (50, 51). In order to investigate the relationship between ATM, ATR and DNA-PK and Nol12 function, HCT WT cells transfected with siSCR or siNOL12 were grown in the presence of specific inhibitors of ATM (KU55933), ATR (VE822) and DNA-PK (NU7441) (52-54). Cells were assayed for apoptosis 24h, 36h and 48h post-knockdown as previously described. While neither KU55933 nor NU7441 affected the induction of apoptosis following NOL12 kd, treatment with VE822 reduced the apoptotic response to that of background (i.e. siSCR + VE822; Figure 6b). Dose-response analyses revealed that VE822 inhibited NOL12 kd-dependent apoptosis with an IC_50_ of 92.2nM (R^2^ = 0.982), 4.9-fold higher than the published IC_50_, while KU55933, for comparison, inhibited apoptosis by only 38% at the highest concentration tested (20μM, 1550-fold over the published IC_50_). Next we tested the relation of NOL12 kd-induced apoptosis to Chk1 and Chk2, the kinases downstream of ATM and ATR. Surprisingly, treatment with the Chk1-inhibitor ChiR124 did not reduce the apoptotic response upon NOL12 kd (Figure 6c, middle), while treatment with the Chk2-inhibitor NSC109555 led to a mild reduction (Figure 6c, right) (55, 56). These results suggest that, while NOL12 kd-induced apoptosis requires signaling via ATR, it may do so independently of Chk1, despite leading to its phosphorylation; instead, apoptosis upon loss of Nol12 appears to be dependent on signaling via Chk2. Importantly, knockdown of the ribosomal protein gene, RPS6, did not result in an apoptotic response and was insensitive to VE822 treatment, confirming that the NOL12 kd-related apoptotic response is unlikely the result of ribosomal stress (Figure 6d). Overall, these data suggests that Nol12 may function in an ATR-induced DNA damage response, which is supported by a lack of co-localization of Nol12 with the ATM-activated DNA double-strand break marker 53BP1 in HCT WT cells, both in absence and presence of 1μg/mL Actinomycin D (ActD), which intercalates into DNA and induces double-strand breaks (data not shown) (57).

Given that the cell cycle arrest arising from NOL12 kd was resistant to inhibition of nucleolar stress pathways, we tested whether it too was dependent on DNA damage signaling. To this end, we carried out cell cycle profiling experiments in the presence of caffeine, a PIKK kinase inhibitor that targets both ATM and ATR, and the DNA-PK inhibitor NU7441 (58). Consistent with observations described above, the <2n cell cycle population (consisting of apoptotic/necrotic cells and cell debris) was significantly decreased in the presence of caffeine but not NU7441 (Figure 6e), as was the total apoptotic cell count (Supplementary Figure 5e). However, the induction of G_1_/S arrest was unchanged in the presence of either caffeine or NU7441 (Figure 6e), suggesting that the cell cycle arrest and apoptotic phenotypes of NOL12 kd arise by different mechanisms and, possibly, different upstream events.

### Loss of Nol12 leads to increased DNA oxidation

The activation of the ATR-Chk1/Chk2 DNA damage pathway and ATR-dependent apoptosis by NOL12 kd led us to re-examine the interactome of PrA-Nol12 with regards to a possible role on damaged DNA. Indeed, PrA-Nol12 co-isolated numerous proteins implicated in DNA damage response and repair pathways. In particular, PRKDC, the catalytic component of DNA-PK, and Dh×9, a multifunctional nuclear helicase that resolves R‐ and D-loops and is excluded from the nucleolus in most cells, were both coisolated in high quantities in all tested conditions (23). Nol12 also co-isolated high amounts of NONO and SfpQ (Figures 1a and 7a), two mRNA processing factors and paraspeckle components that are recruited to sites of DNA damage, and co-localized with SfpQ (Figure 2b) (59, 60). Association of Nol12 with DNA-PK, Dh×9 and NONO was not RNA dependent, suggesting either a DNA-related role or association with these proteins prior to assembly on target RNA; however, with the exception of Dh×9, most associations in this group, including DNA-PK and NONO, were salt-sensitive, suggesting transient or labile interactions. Interestingly, while Nol12 partially co-localized with SfpQ to paraspeckles, upon treatment with ActD at a concentration that specifically inhibits RNA Pol I but not RNA Pol II (0.05 μg/mL, 3 h) inducing to the formation of nucleolar caps, unlike SfpQ, Nol12 did not retreat into nucleolar caps, but instead formed a peripheral nucleolar ring while still maintaining its presence within nucleoplasmic foci and the cytoplasm (Figure 7b). Nol12 also co-localized with Dh×9, and overlap of foci was increased upon ActD treatment at concentrations known to affect overall genomic stability (1 μg/mL, 3 h), suggesting, in line with our proteomic data, a potential co-function for Nol12 in genome instability management (Figure 7c).

**Figure 7.**
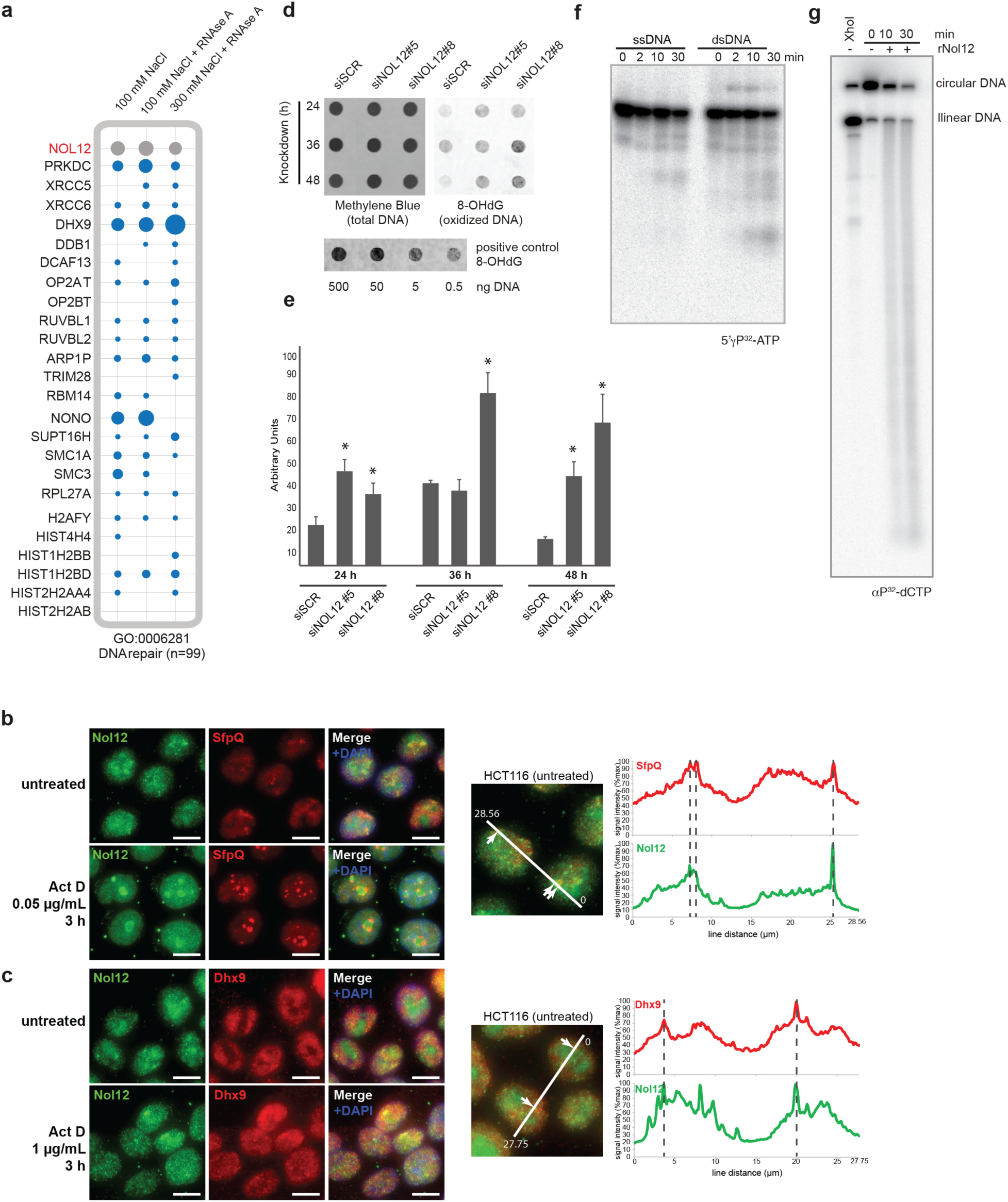
Nol12 associates with DDR factors and is required for suppression of oxidative DNA damage. (a) PrA-Nol12 associated proteins identified by MS analysis with a “DNA repair” gene ontology annotation (GO:0006281). Dots represent semi-quantitative analyses. (b, c) Endogenous Nol12 localization in HCT 116 WT cells was compared to that of the paraspeckle component SfpQ (b, red) and to that of the RNA/DNA DEAD-box helicase Dh×9 (c-red; arrows) in untreated and actinomycin D-treated cells. Images are representative of three independent experiments. Scale bar = 10 μιμ. In each case HCT116 WT cells were grown on poly-L-lysine-coated coverslips for 48h, with or without treatment actinomycin D (either 0.05μg/ml (b) or μg/ml (c), 3h) prior to fixation, imaging and trace analysis. Linear profiles of fluorescence density were created from representative images of each co-immunofluorescence experiment and normalized against the maximum observed fluorescence along the line segment. Arrows/dotted lines indicate the location of local fluorescence maxima in the marker channel, and colocalization was analysed using the Fiji software Coloc2 module. (d) DNA purified from HCT WT cells, siRNA-treated as indicated, was spotted onto a nylon membrane. Oxidized DNA was detected using an anti-8-OHdG antibody, and (e) quantified densitometrically. Data represent mean±s.d. for three independent experiments; * p<0.05 vs. siSCR treatment by Student’s two-tailed t-test. (f) *In vitro* 5′-radiolabeled ssDNA, dsDNA (yP^32^-ATP) or (g) circular dsDNA (αP^32^-dCTP) was incubated for the indicated times with rNol12 and resolved by denaturing polyacrylamide electrophoresis. Images are representative of three independent experiments.

Given the above data, we assayed whether loss of Nol12 was able to accumulate DNA damage sufficient to activate the ATR pathway. Total cellular DNA from HCT WT cells treated with control siSCR, or two different NOL12 siRNAs (siNOL12 #5 and #8) was probed by southwestern blotting with an antibody against 8-hydroxydeoxyguanosine (8-OHdG), a form of oxidative DNA damage shown to predominantly activate ATR-associated DDR (49). Strikingly, 8-OHdG was significantly accumulated in cellular DNA 24-36h after Nol12 depletion (Figure 7d, e), suggesting that Nol12 may be required for protection against 8-OHdG formation in this cell line. This induction of oxidative stress is in accordance with the observed activation of Mdm2^pS166^ (Figure 5d, g), which accumulates in an MEK-ERK-dependent manner upon H_2_O_2_ treatment (48). As this data implies a possible role for Nol12 in protection from DNA damage or in DNA repair, we tested whether the protein, in addition to possessing RNA endonuclease activity, was also able to metabolize DNA substrates *in vitro*. Similar to RNA substrates, incubation of rNol12 with end-labeled, linear ssDNA and dsDNA substrates resulted in the appearance of labeled products of different lengths while no accumulation of radio-labeled nucleotide was observed, consistent with endonucleolytic cleavage of the substrate (Figure 7f). In addition, incubation of rNol12 with ligated, circular radiolabeled dsDNA resulted in linearization of the circular DNA, producing a linear ssDNA by endonucleolytic cleavage and a smear of smaller products over the course of 30 minutes (Figure 7g). These data support an endonucleolytic activity for Nol12 on DNA, identifying it as a RNA/DNA endonuclease *in vitro* with a role in the prevention or repair of DNA oxidation *in vivo*.

## DISCUSSION

We show here that the human protein Nol12 is a RNA/DNA endonuclease required for both ribosome maturation and genome maintenance in higher eukaryotes. Nol12 is part of early 90S and pre-60S ribosomal subunits *in vivo*, where it is required for efficient separation of the large and small subunit precursors via cleavage at site 2. Knockdown of NOL12 impairs cellular proliferation by inducing a G_1_/S cell cycle arrest, but does so outside of a nucleolar stress response and in a p53 and c-myc-independent manner. In addition, Nol12 has a putative role in the prevention and/or resolution of DNA oxidation as loss of the protein leads to an increase in oxidized DNA levels followed by a rapid activation of the DNA damage response kinase ATR and apoptotic response. Moreover, Nol12 interacts with a number of chromatin-associated factors *in vivo*, in particular proteins implicated in DNA damage repair, and colocalizes with paraspeckle component SfpQ, and the RNA/DNA DEAD-box helicase Dh×9, involved in genome instability prevention, suggesting further roles in genome maintenance. Nol12 is also associated with RNA metabolism pathways outside of ribosome biogenesis, namely in P-bodies, which is supported by both proteomic data and its co-localization with Dcp1a.

### Nol12, the nucleolus, ribosome biogenesis and cell growth

Redundant ribosome maturation pathways are considered vital to sustain ribosome levels and, in higher eukaryotes, this link is emphasized by the fact that loss of most ribosome biogenesis factors leads to a cell cycle arrest in G_1_/S, which is always preceded by the appearance of rRNA processing defects; however, while the G_1_/S arrest is part of a nucleolar stress response, it rarely leads to apoptosis in a majority of cell types (37). Our characterization of Nol12’s role in pre-rRNA processing revealed the protein to be required for cleavage at site 2 as all immediate precursors resulting from cleavage at this site - 30S and 21S, but mostly 32.5S - are significantly reduced in the absence of Nol12. Given its *in vitro* endonuclease activity our data suggests that Nol12 may be the as yet unidentified site 2 endonuclease. In its absence the pathway is rerouted through site E as indicated by the increase in 36S and subsequent Xrn2-processed intermediates (36S-C), presumably to ensure ribosome production, which is supported by the moderate effect on mature rRNA levels. A role for Nol12 as the site 2 endonuclease is strongly supported by the Nol12 pre-ribosome interactome, in particular, the significant enrichment of Utp10, Rrp5, Ftsj3, Gtpbp4, and Rsl1D1 (Supplementary Table 3). The association with a high number of 90S factors suggests an early recruitment of Nol12 to pre-rRNA as Utp10 is a t-UTP factor that was suggested to be one of the earliest proteins to associate with the 47S precursor (61). The RNA binding protein Rrp5 was recently shown to be important for both cleavage at site 2 and processing from site 2 to E by Rrp6 (4), while loss of Ftsj3 delayed processing at site 2, shifting events to the minor pathway via site E (62). It is conceivable that these proteins either aid in the recruitment of Nol12 to pre-ribosomes, or facilitate Nol12’s function at site 2, possibly even through direct interaction. Nothing is known about the roles of Rsl1D1 and Gtpbp4 in pre-rRNA processing, yet their depletion results in a similar pattern of precursors than those observed upon loss of Nol12, suggesting a function in or around the same processing events (37). Rcl1, the endonuclease responsible for cleavage at site E, and Xrn2 were co-isolated at very low levels, supporting the notion that they function along a different yet parallel pathway than Nol12 (4). Inconclusive results of previous studies on Nol12’s role in ITS1 processing were most likely due to the fact that processing events were surveyed 60h after siRNA transfection, at which point significant levels of apoptosis can be observed in Nol12-depleted cells (4).

However, besides rerouting of processing events, loss of Nol12 also elicits a strong G_1_/S arrest independent of both p53 and c-myc signaling, and outside a typical nucleolar stress response. Moreover, the scale of the arrest is markedly more severe compared, for example, to that observed upon Xrn2 depletion, while decrease in mature rRNAs, and subsequently ribosomes, is much less severe to knockdowns of other maturation factors, or to warrant the strong and rapid onset of apoptosis in the absence of Nol12 (4, 37, 63). It is therefore likely that Nol12 has other essential cellular functions outside the nucleolus.

### A role for Nol12 in the maintenance of genome integrity

In *Drosophila*, kd of the NOL12-homologue *Viriato* resulted in loss of cell proliferation and apoptosis (20, 64). Oxidative stress, and in particular the production of reactive oxygen species (ROS), is a known contributor to cellular damage, including oxidative DNA damage, and, if not repaired, leads to subsequent cell death (49). The accumulation of 8-OHdG after 24-36h in Nol12-depleted cells is likely the cause for the observed apoptotic response 36-48h post-siRNA treatment. This notion is supported by the activation of the ATR-Chk1/2 DDR pathway and phosphorylation of Chk1^pS345^, Mdm2^pS166^ and Chk2^pT68^ observed after 24h, and the decrease in p53^pS15^ levels at 48h, most likely allowing for Mdm2^pS166^-triggered degradation of p53 (48, 49). Moreover, this rapid activation of the DDR pathway suggests this to be a direct cause upon loss of Nol12. ATR-Chk1-mediated apoptosis has previously been observed upon replicative stress via ribonucleotide reductase inhibition, interstrand cross-links, oxidative or base damage of DNA, amongst others (50). Moreover, the abolition of apoptosis, but not G_1_/S arrest, upon ATR inhibition fully supports an oxidative stress induced apoptosis, and further suggests that these two responses, G_1_/S arrest and cell death, likely arise from different primary stimuli. The observed interactions of Nol12 with DNA damage factors, its localization to transcription-independent nucleoplasmic granules, co-localization with Dh×9, and its ability to endonucleolytically cleave dsDNA all raise the possibility that Nol12 may act directly on damaged DNA. Similarly, Nol12 could either support the resolution of sites of oxidized DNA and/or suppress oxidized DNA formation; loss of either activity would trigger the ATR pathway. Alternatively, the cause of increased DNA oxidation could also be due to the increased generation of ROS, possibly through perturbation of mitochondrial function. The Nol12 interactome contains a significant number of proteins found in the mitochondria (Supplementary Table 3). In particular, a marked enrichment is seen for protein components of the mitochondrial ribosome (MRPLs/MRPSs; (65)) and of proteins found in the mitochondrial nucleoid, the mitochondrial DNA-protein complex containing the mitochondrial genome and proteins required for DNA replication, stability, transcription and early rRNA processing (66). This indicates the possibility that Nol12 may possess functions in DNA repair or ribosome biogenesis within the mitochondria. Loss of these functions would lead to mitochondrial dysfunction and, ultimately, increased intracellular ROS and apoptosis (67). This notion is supported by recent findings of Nol12 interacting with superoxide dismutase 2 (SOD2) (68), an enzyme that catalyzes ROS in mitochondria, limiting their detrimental effects in an anti-apoptotic role against oxidative stress, as well as the neuron-expressed cytosolic protein p33Monox, inhibitor of APP and Bcl2 phosphorylation (24), and APP itself (69). Taken together this may suggest a role for Nol12 in prevention of oxidative damage in neurons. Finally, Nol12 may also act on oxidized RNA, and to date no known definitive enzyme for the repair of oxidized RNA has been identified (18); however, such function remains to be determined.

### Nol12, a nuclease with diverse cellular roles

Multifunctional nucleases are common in many organisms. In higher eukaryotes, the multifunctional RNA exonucleases Xrn2 and hRrp6, were recently shown to aid in the resolution of RNA:DNA hybrids and repair of DNA double-strand breaks, respectively, expanding their functions beyond the RNA realm (13, 14); this function of Xrn2 is also the most likely cause for the phosphorylation H2A.X^pS139^ observed in its absence. The endonuclease Dicer was shown to process both miRNAs and siRNAs, while also functioning during DNA damage repair (8, 15), and the human AP endonuclease 1 (APE1) was previously characterized as a multifunctional enzyme: a major component of the DNA base-excision repair pathway and responsible for decay of *c-myc* mRNA (8). The co-isolation of components from diverse cellular RNA/DNA homeostasis niches, its localization to different subcellular compartments as well as its RNA and DNA endonuclease activity supports the idea of Nol12 as another multifunctional nuclease. Nol12 has also been identified in several recent high-throughput screens for mRNA-binding proteins (70, 71), and, here, we found the protein was localized to GW/P-bodies under normal growth conditions as well as enriched upon arsenite stress, suggesting a putative role in mRNA metabolism. Moreover, Nol12’s association and co-localization with paraspeckle components suggests a potential function within these bodies, either in the cleavage of the 566 adenosine-to-inosine edited Alu-repeat containing human RNAs, as no endonuclease for this function has been identified to date, or during DNA damage (72, 73). Both possibilities will be investigated in the future.

Being a broad-specificity RNA/DNA nuclease, Nol12’s multifunctionality is likely to be conveyed through either different cofactors such as helicases, which would also explain its low *in vitro* nuclease activity, and/or posttranslational modifications (PTMs). A casein kinase II target site has been identified at S134 within both Nol12 and its mouse homologue Nop25, and both the CKII subunit CSNK2A1 and the phosphatase PP1A are part of the Nol12 proteome (Supplementary Table 3). Moreover, several helicases were found significantly enriched with Nol12, including Ddx21, which is involved in both transcription and rRNA processing (74), the GW/P-body proteins Ddx5 and Ddx17 (75), and the multifunctional RNA/DNA helicase Dh×9, implicated in the resolution of numerous aberrant DNA structures including triple-helices, G-quadruplexes, D-loops and, most significantly, RNA:DNA hybrids (76). Dh×9 has also been shown to co-localize with H2A.X^pS139^ and DNA-PK in RNA-containing granules upon exposure of cells to low-dose actinomycin D and to protect Eμ-*myc*/Bcl-2 lymphoma cells against an ATR-dependent apoptotic response (77), suggesting it may be required for the resolution of stalled RNA polymerases and/or the resulting RNA:DNA hybrids. Two other Nol12 interactors, SUPT16H and SSRP1, have also been implicated in R-loop resolution in humans (78). Hence, a role for Nol12 in the resolution of unusual DNA structures or RNA:DNA hybrids in concert with Dh×9 would provide another attractive explanation for the activation of an ATR-dependent apoptotic response following Nol12 depletion. The recent identification of RNA processing proteins in the DNA damage response by a number of screens, speaks for a functional juncture between RNA metabolism and DNA repair pathways (19). Overall, the putative multifunctionality of Nol12 presents one such protein at the intersection of RNA metabolism and DNA maintenance and repair, and it will be interesting to explore the mechanisms of its different functions in the future.

## ACKNOWLEDGMENTS

The authors thank Drs. J. Archambault, N. Francis, E. Lecuyer, F. Robert, N. Watkins, and D. Zenklusen for gifts of reagents used in this study, Dr. C. Denicourt for cell lines, S. Rahman and P. Raymond for assistance with microscopy, D. Faubert of the IRCM Proteomics platform, and members of the Oeffinger and Zenklusen labs for critical reading.

## FUNDING

This work was supported by operating grants from the Canadian Institutes of Health Research (MOP 106628-MO); the Natural Sciences and Engineering Research Council of Canada (RGPIN 386315-MO); and Fonds de recherche du Québec – Santé (PR2I). M.O. holds a CIHR New Investigator award and Fonds de recherche du Québec - Santé Chercheur Boursier Junior I. Funding for open access charge: Natural Sciences and Engineering Research Council of Canada.

